# Rare variants imputation in admixed populations: Comparison across reference panels and bioinformatics tools

**DOI:** 10.1101/494229

**Authors:** Sanjeev Sariya, Joseph Lee, Richard Mayeux, Badri N. Vardarajan, Dolly Reyes-Dumeyer, Jennifer Manly, Adam Brickman, Rafael Lantigua, Martin Medrano, Ivonne Z. Jimenez-Velazquez, Giuseppe Tosto

## Abstract

**Background:** Imputation has become a standard approach in genome-wide association studies (GWAS) to infer *in silico* untyped markers. Although feasibility for common variants imputation is well established, we aimed to assess rare and ultra-rare variants’ imputation in an admixed Caribbean Hispanic population (CH).

**Methods:** We evaluated imputation accuracy in CH (N = 1,000), focusing on rare (0.1% ≤minor allele frequency (MAF) ≤ 1%) and ultra-rare (MAF < 0.1%) variants. We used two reference panels, the Haplotype Reference Consortium (HRC; N = 27,165) and 1000 Genome Project (1000G phase 3; N = 2,504) and multiple phasing (SHAPEIT, Eagle2) and imputation algorithms (IMPUTE2, MACH-Admix). To assess imputation quality, we reported: a) high-quality variant counts according to imputation tools’ internal indexes (e.g. IMPUTE2 “Info”≥80%). b) Wilcoxon Signed-Rank Test comparing imputation quality for genotyped variants that were masked and imputed; c) Cohen’s kappa coefficient to test agreement between imputed and whole-exome sequencing (WES) variants; d) imputation of G206A mutation in the *PSEN1* (ultra-rare in the general population an more frequent in CH) followed by confirmation genotyping. We also tested ancestry proportion (European, African and Native American) against WES-imputation mismatches in a Poisson regression fashion.

**Results:** SHAPEIT2 retrieved higher percentage of imputed high-quality variants than Eagle2 (rare: 51.02% vs. 48.60%; ultra-rare 0.66% vs 0.65%, Wilcoxon p-value < 0.001). SHAPEIT-IMPUTE2 employing HRC outperformed 1000G (64.50% vs. 59.17%; 1.69% vs 0.75% for high-quality rare and ultra-rare variants, respectively; Wilcoxon p-value < 0.001). SHAPEIT-IMPUTE2 outperformed MaCH-Admix. Compared to 1000G, HRC-imputation retrieved a higher number of high-quality rare and ultra-rare variants, despite showing lower agreement between imputed and WES variants (e.g. rare: 98.86% for HRC vs. 99.02% for 1000G). High Kappa (*K* = 0.99) was observed for both reference panels. Twelve G206A mutation carriers were imputed and all validated by confirmation genotyping. African ancestry was associated with higher imputation errors for uncommon and rare variants (p-value < 1e-05).

**Conclusion:** Reference panels with larger numbers of haplotypes can improve imputation quality for rare and ultra-rare variants in admixed populations such as CH. Ethnic composition is an important predictor of imputation accuracy, with higher African ancestry associated with poorer imputation accuracy.

## Introduction

Genome-wide association studies (GWASs) are a major tool to identify common variants associated with complex diseases. GWAS can include 550K to over 2M Single Nucleotide Polymorphisms (SNPs) (Ha et al., 2014) to cover the human genome evenly. Although GWAS has shown to be a robust method to identify disease loci of interest, they rarely point to a causal coding variant. In fact, microarray SNP chips for GWAS are optimally designed to uncover common variants, often associated with small effect sizes mostly located in intronic and intergenic regions. The focus of genetic investigations has since shifted toward rarer alleles with larger effect sizes (Gibson, 2012). With the changing paradigm, imputation of rare variants has become an important topic to enhance the genome coverage in GWAS. Imputation is a process of inferring untyped SNP markers in the discovery population by using densely typed SNPs in external reference panel(s). These ‘*in silico*’ markers increase the coverage of association tests while conducting genome-wide association analysis. In addition, large number of SNPs facilitate meta-analysis when merging data from different study cohorts.

The quality of imputation essentially depends on two parameters: available reference datasets and algorithms that employ those reference datasets. Previous studies have shown that imputation quality depends on how well reference panels reflect the study population. To respond to the needs, the 1000 Genome project (1000G), now in its third phase release, has proven to be one of the most frequently used reference panels (Genomes Project et al., 2015). Using these composite reference panels, a number of studies (Pei et al., 2010; Howie et al., 2012; Verma et al., 2014; Liu et al., 2015) have compared imputation accuracy using different imputation tools and algorithms, although the results are equivocal. Few studies (Browning and Browning, 2009; Zheng et al., 2012; Zheng et al., 2015) assessed the impact of reference panel size and input data’s features - such as density of SNPs - to impute rare variants, suggesting larger size of reference panels work better. Surakka and colleagues (Surakka et al., 2016) assessed accuracy of imputed SNPs by evaluating rate of false polymorphisms in a Finnish population using global reference panels – Haplotype Reference Consortium (HRC) release 1, 1000G phase 1 and a local reference panel. They concluded that higher false positive rate was observed in imputation from global reference panels compared to imputation performed using a local panel. Other studies (Huang et al., 2015; Das et al., 2016) found imputation accuracy increases with higher number of haplotypes, specifically for variants with MAF ≤ 0.5%. For Hispanic populations, Nelson and colleagues (Nelson et al., 2016) compared imputation performances with 1000G phase 1 (N = 1,092) vs. 1000G phase 3 (N = 2,504), concluding that phase 3 improved accuracy for variants with MAF <1% by. Further, Nagy and colleagues (Nagy et al., 2017) showed that HRC reference panel provides new insight for novel variants particularly for rare variants in a family-based Scottish study cohort. Aforementioned studies highlighted the need of a larger sized reference panel to improve imputation quality. Herzig and colleagues (Herzig et al., 2018) assessed tools for haplotype phasing and their impact on imputation in a population isolate of Campora in southern Italy, and showed that SHAPEIT2, SHAPEIT3 and EAGLE2 were highly accurate in phasing; MINIMAC3, IMPUTE4 and IMPUTE2 were found to be reliable for imputation. Roshyara and colleagues (Roshyara et al., 2014) compared MaCH-Admix, IMPUTE2, MACH, MACH-Minimac in different ethnicities by evaluating accuracy of correctly imputed SNPs; MaCH-Minimac outperformed SHAPEIT-IMPUTE2 in subsamples of different ethnic groups. These studies demonstrated how employed imputation algorithm determines quality of inferred SNPs.

However, no study to our knowledge has evaluated reference panels in tandem with different imputation algorithms to assess imputation quality of inferred SNPs based on MAF in a three-way admixed population. Based on these findings, we assessed imputation quality, focusing on rare and ultra-rare variants, in a large dataset of Caribbean Hispanics (CH) leveraging available GWAS and sequencing data available for our cohort.

## Methods

We will refer SNPs with MAF between 1-5% as “uncommon,” 0.1-1% as “rare,” and ≤0.1% as “ultra-rare”. We considered SNPs with IMPUTE-Info metric ≥0.40 as “good-quality” and ≥0.80 as “high-quality”.

### GWAS samples and genotyping

We selected randomly 1,000 Caribbean Hispanics as part of an original genotyped cohort of 3,138 individuals: genotyped data can be downloaded at dbGaP Study Accession: phs000496.v1.p1. 719 individuals were derived from Estudio Familiar Investigar Genetica de Alzheimer (EFIGA), a study of familial LOAD; and 281 individuals from the multiethnic longitudinal cohort, Washington Heights, Inwood, Columbia Aging Project (WHICAP). The information on study design, recruitment and GWAS methods for the EFIGA and WHICAP study was previously described in Tosto, G., et al (Tosto et al., 2015).

### GWAS quality control (QC)

Genotyped data underwent quality control using PLINK (v1.90b4.9 64-bit) (Purcell et al., 2007). Briefly, we excluded SNPs with missing rate ≥5% followed by exclusion of SNPs with MAF ≤ 1%. We then removed SNPs with P-value < 1e-6 for Hardy-Weinberg Equilibrium. Samples with missing call rate ≥5% were excluded from analysis.

### Global Ancestry estimation and selection of “true Hispanics”

Prior to imputation, we estimated global ancestry using the ADMIXTURE (v.1.3.0) software (Alexander et al., 2009; Zhou et al., 2011). We conducted supervised admixture analyses using three reference populations: African Yoruba (YRI) and non-Hispanic white of European Ancestry (CEU) from the HAPMAP project as representative of African and European ancestral populations; and eight Surui, 21 Maya, 14 Karitiana, 14 Pima and seven Colombian individuals from the Human Genome Diversity Project (HGDP) were used to represent native American ancestry (Li et al., 2008). We used ∼80,000 autosomal SNPs that were: I) genotyped in all three datasets (Caribbean Hispanics, 1000G and HGDP); II) common (i.e. MAF >5%); and III) in linkage equilibrium. Supervised admixture analyses with the three reference populations (YRI, CEU, and Native Americans) revealed that European lineage accounted for most of the ancestral origins (59%), followed by African (33%) and native American ancestry (8%). We then selected only individuals with at least 1% of all three ancestral populations.

### Reference panels

HRC reference panel contained over 39M SNPs from 27,165 individuals who participated in 17 different studies (Table 1). The data were downloaded from the Wellcome Trust Sanger Institute (WTSI). 1000G phase 3 reference panel contained over 81M SNPs from 2,504 individuals (https://mathgen.stats.ox.ac.uk/impute/1000GP_Phase3.tgz). It includes 26 ethnic groups, with most variants rare, approximately 64 million had MAF <0.5%; approximately 12 million had a MAF between 0.5% and 5%; and approximately 8 million have MAF >5%. In order to perform imputation with MaCH-Admix, 1000G Phase 3 pre-formatted data were downloaded from ftp://yunlianon:anon@rc-ns- ftp.its.unc.edu/ALL.phase3_v5.shapeit2_mvncall_integrated.noSingleton.tgz that contained over 47M SNPs. The subsequent analyses were restricted to autosomal chromosomes, only.

**Table 1:** SNP counts in HRC and 1000G reference panel.

| Reference Panel | Individuals | Autosomal variants | Bi-allelic SNPs | Multi-allelic SNPs |
| --- | --- | --- | --- | --- |
| **1000G Phase 3** | 2,504 | 81,706,022 | 77,818,332 | 3,887,690 |
| **HRC** | 27,165* | 39,131,600 | 39,131,600 | NA |

\*For Chromosome 1, the number of individuals were 22,691

### Phasing and Imputation Procedures

We compared SHAPEIT2 (Delaneau et al., 2013) and Eagle2 (Loh et al., 2016) by phasing and then imputing (see next section) a single chromosome (Chromosome 21), using both reference panels. We refer to SHAPEIT2 as SHAPEIT when used in tandem with IMPUTE2 for the remainder of paper.

Imputation was carried out using two bioinformatics tools: IMPUTE2 (Howie et al., 2009) and MaCH-Admix (Liu et al., 2013). For both, imputation quality ranged from 0 to 1, with 0 indicating complete uncertainty in imputed genotypes, and 1 indicating no uncertainty in imputed genotypes.

### IMPUTE2 (version 2.3.2)

IMPUTE2 uses an MCMC algorithm to integrate over the space of possible phase reconstructions for genotypes data. We conducted imputation in non-overlapping 1MB chunk regions; chunk coordinates were specified using the “–*int*” option. Other options were used with default parameters (**Supplementary section 1**). Briefly, we used a default 250KB buffer region to avoid quality deterioration on the ends of chunk region. “-Ne” value as 2000 suggested for robust imputation which scales linkage disequilibrium and recombination error rate. <u>MaCH-Admix</u>. We used MaCH-Admix because it uses a method based on IBS matching in a piecewise manner. The method breaks genomic region under investigation into small pieces and finds reference haplotypes that best represent every small piece, for each target individual separately. MaCH-Admix imputes in three steps: phasing, estimation of model parameter that includes error rare and recombination rate and lastly, haplotype-based imputation. MaCH-Admix (version Beta 2.0.185) was run on default parameters of 30 rounds, 100 states (– autoFlip flag). Details can be found in supplementary file (**section 1**). We initially compared performance between MaCH-Admix and IMPUTE2 using the 1000G reference panel for Chromosome 21 only. We then proceeded to impute all remaining chromosomes with the tool that performed better.

### Imputation Performance Metrics

IMPUTE2 uses “Info” parameter to report imputation quality that measures relative statistical information about SNP allele frequency from imputed data. It reflects the information in imputed genotypes relative to the information if only the allele frequency were known. “Info” metric is used to filter poorly imputed SNPs from IMPUTE2 and is reported for all imputed SNPs. In addition, IMPUTE2 uses an internal metric known as R^2^, reported for genotyped SNPs only: it measures squared correlation between genotyped SNPs and the same SNPs that have been first masked internally and then imputed. MaCH-Admix uses *Rsq* to report imputation quality. The R^2^ metric is also known as variance ratio, calculated as proportion of empirically observed variance (based on the imputation) to the expected binomial variance p(1−p), where p is the minor allele frequency. A threshold of 0.30 is recommended to filter out poorly imputed SNPs.

Despite quality measures from IMPUTE2 and MaCH-Admix being highly correlated (Marchini and Howie, 2010), we calculated a *r2hat* score to generate a single common metric to assess imputation quality across the software (Hancock et al., 2012) (v109, http://www.unc.edu/∼yunmli/tgz/r2_hat.v109.tgz).

We compared performance of MaCH-Admix and SHAPEIT-IMPUTE2 by: a) Reporting raw SNP counts based on quality (MaCH-Admix “Rsq” and IMPUTE2 “Info”); b) Comparing *r2hat* for overlapping imputed SNPs from both tools; c) Conducting a Wilcoxon Signed-Rank Test (R v3.4.2) on *r2hat* value of overlapping SNPs.

We compared performance of Eagle2 and SHAPEIT2 phasing tools in tandem with IMPUTE2 as imputation tools across reference panels by: a) Comparing their respective IMPUTE2 R^2^: b) Conducting a Wilcoxon Signed-Rank Test on R^2^ value; c) Reporting raw counts of imputed SNPs based on IMPUTE2 “Info” metric and stratified by MAF bins (e.g. common, rare, ultra-rare).

In all comparisons, the MAFs are estimated from imputed data according to the reference panel employed. We retained monomorphic SNPs in our analyses for several reasons. A monomorphic SNP in one study might not be monomorphic in other cohorts. This has profound affects, for example, when performing meta-analysis across different studies. In addition, monomorphic SNPs provide information about MAF across studies. Without the information it is difficult to tell, for instance, if a SNP is monomorphic or failed quality control in that study.

### Agreement between Imputed and Sequence data

To further test the quality of imputation - without relying on software’s internal metrics (i.e. “Info” and R^2^) - we calculated genotyped concordance between imputed and WES data using the VCF-compare tool (v0.1.14-12-gcdb80b8) (Danecek et al., 2011). First, we converted posterior probabilities obtained from imputation into genotype data using the PLINK software (v1.90b4.9) by applying a threshold of 0.9 (**supplementary section 1**), such that SNPs that failed on this criterion were left uncalled. For example, an imputed SNP with P(G = 0,1,2)= (0.01,0.9,0.09) would be called as a ‘1’ (heterozygous), whereas an imputed SNP with P(G = 0,1,2) = (0.2, 0.6, 0.2) would be left uncalled. We restricted the comparison to overlapping SNPs between HRC, 1000G reference panels and whole-exome sequencing (WES) data for Chromosome 14 only, on SNPs with 0% missingness (plink –missing flag) in WES data. We also assessed variants’ agreement according to different MAF bins for “high-quality” (“Info” ≥0.8) SNPs. The output resulted in number of variant “mismatches”, i.e. the count of allele not matching between imputed and sequenced variants per individual. To measure interrater reliability we computed Cohen’s kappa coefficient (McHugh, 2012) for both the reference panels against WES data. Kappa coefficient ≤ 0 indicates no agreement, 0.01–0.20 as none to slight, 0.21–0.40 as fair, 0.41– 0.60 as moderate, 0.61–0.80 as substantial, and 0.81–1.00 as almost perfect agreement.

### Effects of Ancestry on Imputation Quality

To assess how ancestry affected imputation quality, we conducted a Poisson regression using R. We used percentage of global ancestry (European (CEU), Native (NAT) and African (YRI) as predictors, and total number of mismatches as the outcome; analyses were restricted to “high-quality” SNPs, only.

### Imputation of G206A Mutation in PSEN1

To evaluate imputation performance of a specific rare variant, we examined a founder mutation, p.Gly206Ala (G206A - rs63750082) in the *PSEN1* gene (PSEN1-G206A) (Athan et al., 2001; Lee et al., 2015). The PSEN1-G206A mutation is a rare variant observed primarily in Puerto Ricans with familial early onset Alzheimer’s disease (EOAD), but it is rare in Puerto Ricans and other populations with late-onset Alzheimer’s disease (LOAD) (Arnold et al., 2013). The mutation was present in the 1000G phase 3 reference panel with an allele frequency of 0.001, but was absent in the HRC reference panel. To verify whether individuals who were found to carry the PSEN1-G206A mutation based on 1000G-imputation, they were genotyped using the KASP genotyping technology by LGC genomics (https://www.lgcgroup.com), which uses allele-specific PCR for SNP calling. Agreement between imputed and genotype data for the PSEN1-G206A mutation was then assessed. We also tested the effect on imputation quality based on different IMPUTE2-parameters settings, more specifically by modifying the chunk size (i.e. 1MB vs. 5 MB).

## Results

### Comparison of Phasing Tools: Eagle2 vs. SHAPEIT2

To select the optimal tool for phasing, we compared SHAPEIT2 with Eagle2 using Chromosome 21 with 13,066 genotyped SNPs by performing subsequent imputation with IMPUTE2 on phased outputs, and using both reference panels. We found SHAPEIT2 better than Eagle2 when evaluated based on mean R^2^ and “Info” metric using either the reference panels. For instance, using the 1000G, we observed higher mean R^2^ for data phased with SHAPEIT2 as compared to Eagle2 (0.92 vs. 0.91; Wilcoxon p-value < 0.001). Similarly, when HRC panel was employed, mean R^2^ of 0.89 was observed for SHAPEIT2 against 0.85 for Eagle2 (Wilcoxon signed-rank test p-value < 0.001).

SNP count comparison details can be found in **Supplementary Table 1-2**. Regardless of the reference panel employed, we observed higher percentage of “high-quality” rare and ultra-rare SNPs for SHAPEIT-IMPUTE2 than Eagle2-IMPUTE2. For instance, 1000G-imputation retrieved 51.02% of “high-quality” rare SNPs using SHAPEIT-IMPUTE2 vs. 48.38% with Eagle2-IMPUTE2. Detailed comparisons for different MAF bins and quality threshold can be found in **Supplementary Section 2**. Nevertheless, we found Eagle2 faster than SHAPEIT2 when computation times were compared; for instance, with HRC Eagle2 was ∼6 times faster than SHAPEIT2 (**Supplementary Table 3**). We therefore imputed the remaining chromosomes on phased output from SHAPEIT2. Comparison of phasing tools by assessing switch error rate was beyond the scope of this paper due to limited resources, for e.g., availability of phased reference panel for an admixed population.

### MaCH-Admix vs. IMPUTE2

We found that SHAPEIT-IMPUTE2 performed better than MaCH-Admix. For Chromosome 21, we imputed 1,104,648 and 646,594 SNPs for SHAPEIT-IMPUTE2 and MaCH-Admix, respectively; 549,091 SNPs were overlapping. For SHAPEIT-IMPUTE2 we observed 446,591 bi-allelic SNPs with “Info” ≥ 0.40, in contrast with 598,943 SNPs with Rsq ≥ 0.30 from MaCH-Admix (**Supplementary Table 4**). SNP counts for different MAF bins based on platform-specific quality index can be found in **Supplementary Table 5**.When the two outputs were compared in terms of *r2hat*, SHAPEIT-IMPUTE2 showed a higheraverage r2hat of 0.62 against 0.36 from MaCH-Admix (Wilcoxon signed-rank test p-value < 0.001). Also, MaCH-Admix was 109 times slower than IMPUTE2. (**Supplementary Table 6),** thus, comparison between different panels using MaCH-Admix were excluded due to limited resources. For the remaining of this manuscript, we focused on imputation employing SHAPEIT-IMPUTE2, only.

### Comparison between HRC and 1000G using SHAPEIT-IMPUTE2

Using SHAPEIT-IMPUTE2, we imputed 81,240,392 and 38,532,090 SNPs across all autosomal chromosomes with 1000G and HRC reference panels, respectively (Table 2).

**Table 2:** Type of imputed SNPs across reference panels.

| Reference Panel | Multi-allelic SNPs |  |  | Bi-allelic SNPs |  |  | Total SNPs |  |  |
| --- | --- | --- | --- | --- | --- | --- | --- | --- | --- |
| | Total SNPs | Info* $\geq 0.40$ (%) | Info $\geq 0.80$ (%) | Total SNPs | Info $\geq 0.40$ (%) | Info $\geq 0.80$ (%) | Total SNPs | Info $\geq 0.40$ (%) | Info $\geq 0.80$ (%) |
| **All SNPs** |  |  |  |  |  |  |  |  |  |
| **1000G** | 3,319,815 | 2,586,342 (77.90) | 2,061,295 (62.09) | 77,920,577 | 31,423,926 (40.32) | 23,468,086 (30.11) | 81,240,392 | 31,423,926 (41.86) | 25,529,381 (31.42) |
| **HRC** | NA | NA | NA | 38,532,090 | 23,436,980 (60.82) | 18,833,790 (48.87) | 38,532,090 | 23,436,980 (60.82) | 18,833,790 (48.79) |
| **SNPs overlapping HRC & 1000G** |  |  |  |  |  |  |  |  |  |
| **1000G** | NA | NA | NA | 30,090,251 | 22,631,112 (75.21) | 18,408,585 (61.17) | 30,090,251 | 22,631,112 (75.21) | 18,408,585 (61.17) |
| **HRC** | NA | NA | NA | 30,090,251 | 22,438,268 (74.56) | 18,395,036 (61.13) | 30,090,251 | 22,438,268 (74.56) | 18,395,036 (61.13) |

Overall, we observed slightly higher mean R^2^ with 1000G than with HRC panel (0.94 vs. 0.92; Wilcoxon p-value < 0.001). Nevertheless, when the analyses were restricted to only “good-” and “high-quality” SNPs, HRC consistently performed better: 60.82% of HRC-imputed SNPs were “good-quality” and 48.87% were “high-quality” (Wilcoxon signed-rank test p-value < 0.001). On the contrary, 40.32% of 1000G imputed SNPs were “good-quality” and 30.11% were “high-quality”.

Further, we evaluated performance for uncommon, rare and ultra-rare SNPs. For “good-” and “high-quality” SNPs, HRC outperformed 1000G. For example, HRC panel produced 62.85% of “high-quality” rare SNPs, whereas 1000G had 53.83% (Table 3). When average imputation “Info” quality was evaluated, HRC-imputation again performed better than with 1000G (Wilcoxon p-value < 0.001) (Figure 1).

**Table 3:** SNP Counts for all Bi-allelic uncommon, rare and ultra-rare SNPs.

| MAF | 1000G |  |  | HRC |  |  |
| --- | --- | --- | --- | --- | --- | --- |
| | Info $\geq 0$ | Info $\geq 0.40$ (%) | Info $\geq 0.80$ (%) | Info $\geq 0$ | Info $\geq 0.40$ (%) | Info $\geq 0.80$ (%) |
| **All SNPs** |  |  |  |  |  |  |
| [1% - 5%] | 6,025,281 | 5,989,223<br>(98.90) | 5,441,982<br>(90.31) | 5,434,996 | 5,421,257<br>(99.84) | 5,061,904 (93.13) |
| [0.1% - 1%) | 20,249,058 | 16,881,286<br>(83.36) | 10,901,789<br>(53.83) | 11,780,671 | 10,931,924<br>(92.79) | 7,404,808<br>(62.85) |
| [0 – 0.1%) | 44,562,205 | 1,490,434<br>(3.34) | 242,717<br>(0.544) | 15,055,433 | 828,256<br>(5.50) | 174,673<br>(1.16) |
| **SNPs overlapping HRC & 1000G** |  |  |  |  |  |  |
| [1% - 5%] | 5,624,956 | 5,604,308<br>(99.63) | 5,148,285<br>(91.52) | 5,396,207 | 5,385,364<br>(99.79) | 5,037,187<br>(93.34) |
| [0.1% - 1%) | 11,875,603 | 10,442,603<br>(87.93) | 7,027,312<br>(59.17) | 10,945,899 | 10,268,136<br>(93.80) | 7,060,908<br>(64.50) |
| [0 – 0.1%) | 6,314,479 | 312,967<br>(4.95) | 47,614<br>(0.75) | 7,519,807 | 560,043<br>(7.44) | 127,423<br>(1.69) |

**Figure 1:**
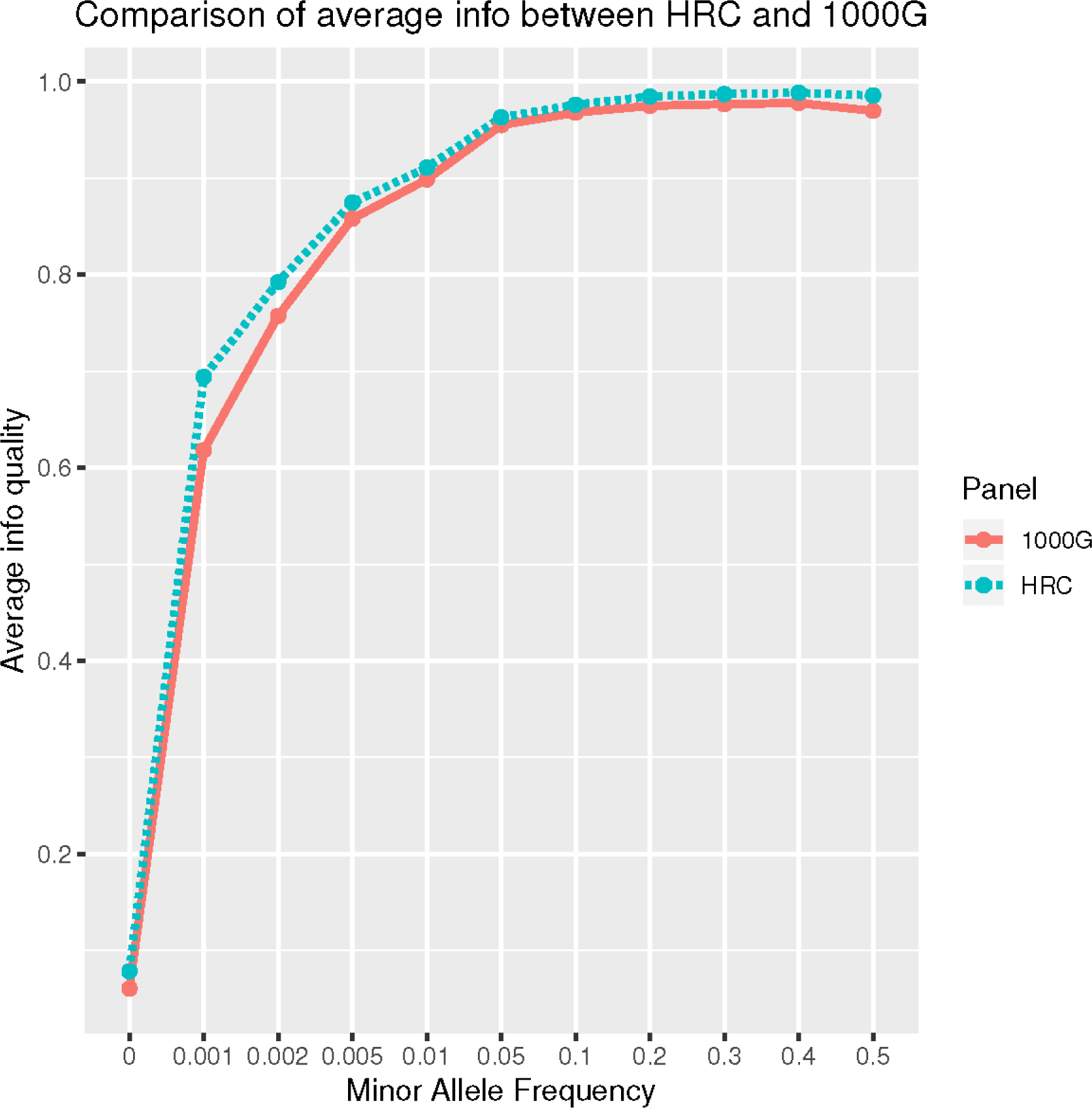
Comparison of average Info quality between HRC and 1000G reference panel for all autosomal chromosomes.

Next, we restricted our analyses to *overlapping* SNPs across the two reference panels only, based on their chromosome and position mapping, reference and non-reference alleles. For “good-”and “high-quality” SNPs, imputation in both panels performed similarly (Table 2). When restricted to uncommon, rare and ultra-rare SNPs, we observed higher percentage of “good-” and “high-quality” SNPs for HRC panel as compared to 1000G reference panel (Table 3). For example, 7.44% of HRC-imputed ultra-rare SNPs were “good-quality” vs. 4.95% with the 1000G. 1.69% of HRC-imputed ultra-rare SNPs were “high-quality” vs. 0.75% with the 1000G. Further, Wilcoxon test on “Info” value of “high-quality” ultra-rare SNPs (2,972) again showed better performances when HRC was employed vs. 1000G (P-value < 0.001). Complete list of counts and percentages across reference panels, MAF bins and quality score can be found in Table 3.

### The case of G206A and the effect of chromosomal chunk size on imputation quality

SNP rs63750082 is absent from HRC panel therefore no imputation was achieved. Using 1000G reference panel, 12 individuals were imputed as G206A carriers. SNP rs63750082 was imputed with an IMPUTE2 “Info” score of 0.48 using 1MB as chromosomal region parameter. When we increased the chunk size to 5MB, IMPUTE-Info score drastically improved to 0.94 (Figure 2). Those patients labeled as mutation-carriers according to imputation were then genotyped: all 12 were confirmed to be G206A carriers, therefore achieving a perfect imputation prediction (100% agreement) for that specific SNP.

**Figure 2:**
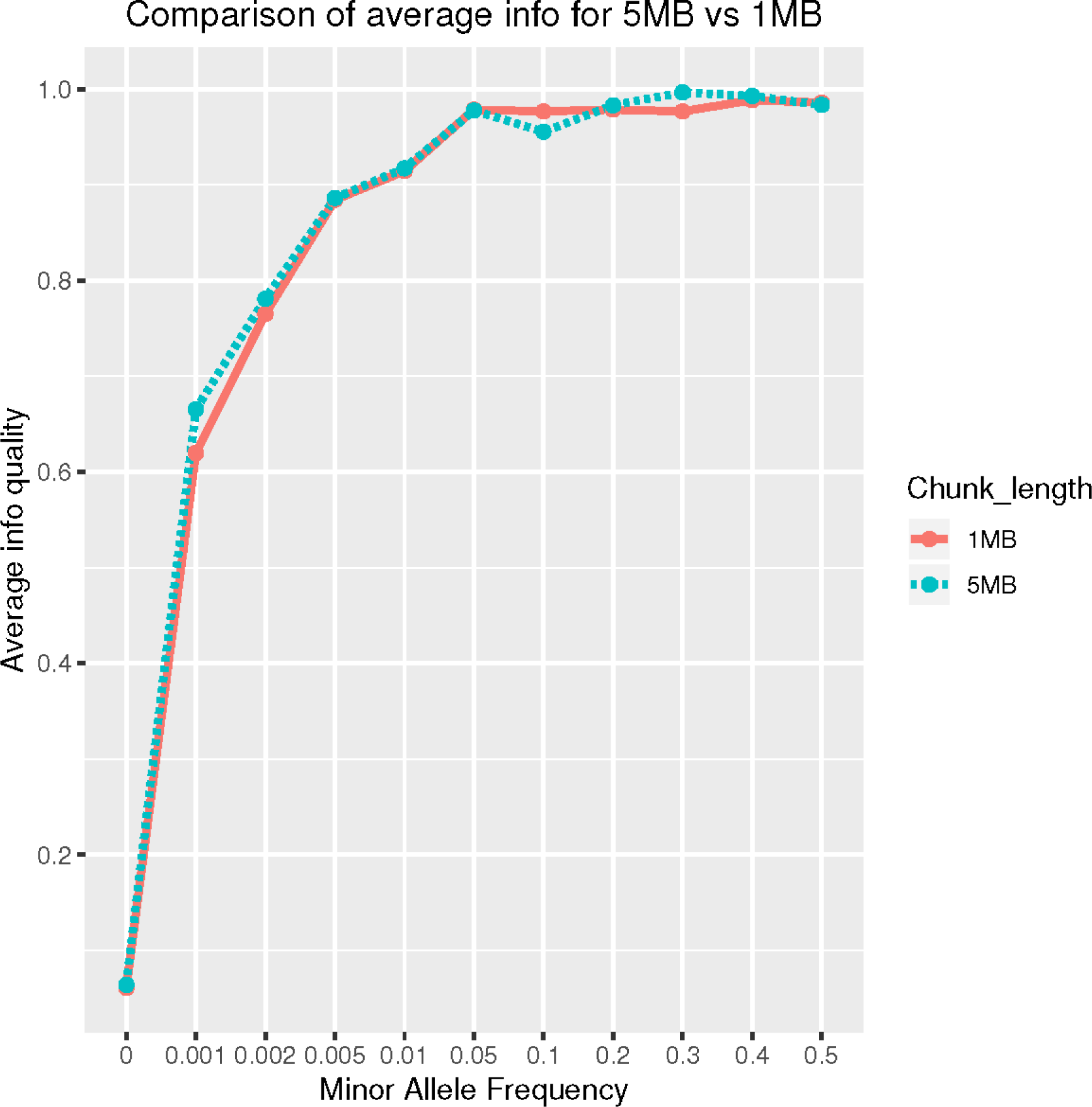
Comparison of average Info on CHR14: 70-75MB (5MB) vs 73-74MB (1MB) region.

### Genotype Concordance and Kappa Coefficient

Out of the 1,000 individuals included in our study, 262 had whole exome sequencing (WES) data available (Raghavan et al., 2018). We had 14,157 overlapping SNPs in WES, HRC and 1000G reference panels with 0% missingness in WES data on Chromosome 14; SNPs imputed with each reference panel were compared against WES data separately. When concordance was evaluated, HRC panel performed slightly poorer, despite showing higher number of “high-quality” variants as compared to 1000G (Table 4). Using 1000G, we observed 3,542 rare and 35 ultra-rare “high-quality” SNPs; across 262 samples, we counted 1,245 ((1,245/(3,542*262))*100 = 0.13%) and 10 (0.10%) mismatches for rare and ultra-rare respectively. Using HRC, we retrieved 3,759 rare and 93 ultra-rare “high-quality” variants; we observed 2,439 (0.24%) and 32 (0.13%) mismatches for rare and ultra-rare variants, respectively. Details about pipeline can be found in **supplementary section 3**. Next, we computed Cohen’s kappa coefficient (*K*) for 14,157 imputed SNPs common in WES and the two reference panels. For both HRC and 1000G-imputation, we observed Kappa (*K*) of ∼0.99 for both rare and ultra-rare “high-quality” variants(Table 4). Details about pipeline can be found in **supplementary section 4**.

**Table 4:**
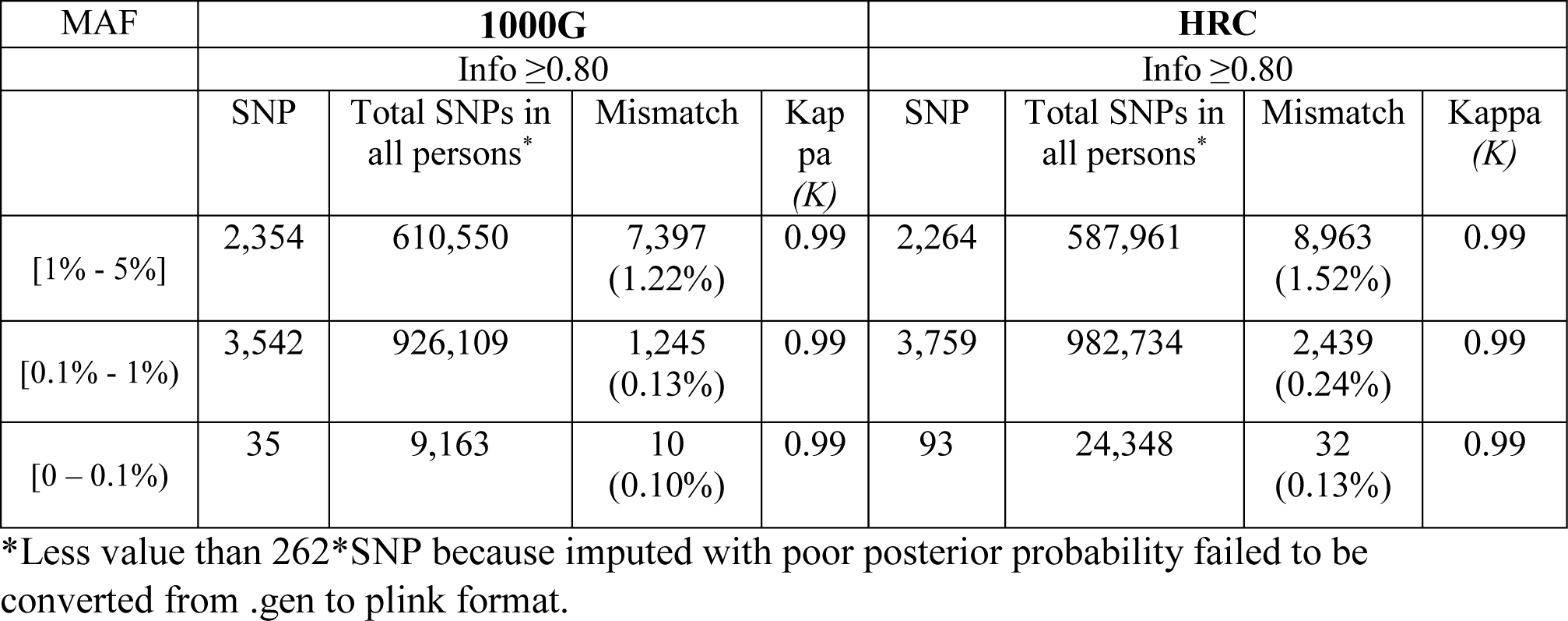
Comparison for mismatch counts and Kappa (*K*) for HRC and 1000G using WES data on Chromosome 14.

### Effects of Ancestry on Imputation Quality

We evaluated the effect of individual ancestral component separately on SNP mismatches for Chromosome 14 on 262 individuals. For both reference panels we found that higher African ancestry (YRI) was associated with higher number of mismatches (**Supplementarytable 7**). For instance, with 1000G reference panel, for rare variants (“Info” ≥0.80), we observed an estimate of 1.46 (P-value < 0.001) for YRI component (indicating that for each unit increase in YRI ancestry, it results in 1.46 additional mismatches). Details on confidence intervals and robust standard errors can be found in **supplementary file (Table 7 and Section 5)**. We did not observe significant effect of ancestry on “high-quality” ultra-rare variants in both panels.

## Discussion

This study examined imputation performances in a cohort Caribbean Hispanics, focusing on uncommon, rare and ultra-rare variant, by comparing different phasing and imputation tools, as well as evaluating the effects of different reference panels. Overall, uncommon and rare variants can be well imputed in this population, characterized by a unique genetic background. Caribbean Hispanics are admixed with 59% of their genetic component from European, 32% African, and 8% Native American ancestry (Tosto et al., 2015). Due to their genetic makeup and unique linkage disequilibrium patterns, admixed populations offer unique opportunity in studying complex diseases. First, disease prevalence varies across ethnic groups (Igartua et al., 2015) and certain admixed populations show higher incidence rates and prevalence (e.g. Alzheimer’s disease, diabetes etc.) or lower ones (e.g. multiple sclerosis). Second, variants that are ethnic-specific may explain a higher prevalence of the disease of interest in admixed groups.

In the present study, we examined multiple parameters of imputation using the Caribbean Hispanics population. First, we found that imputation using SHAPEIT-IMPUTE2 phasing generated better results than Eagle2-IMPUTE2, and SHAPEIT-IMPUTE2 is superior to MaCH-Admix in terms of imputation performances and process time.

Using SHAPEIT-IMPUTE2, 1000G SNPs outnumbered HRC panel because of the higher number of SNPs included in the reference panel itself. However, when we restricted our analyses to overlapping “good-” and “high-quality” SNPs (i.e. those variants that most likely would be included in association analyses), HRC-imputation outperformed 1000G with higher. The superior performance of HRC over 1000G was confirmed also when we focused on uncommon, rare and ultra-rare SNPs only. Our findings confirm data in literature, i.e. reference panels with higher number haplotypes perform better in different scenarios. Additional investigations are needed in order to apply our findings to other admixed and non-admixed populations.

Overall, higher quality of imputation for rare and ultra-rare variants was also confirmed when we tested results against sequencing data. Finally, higher YRI global ancestry was found to significantly impair SNP imputation, suggesting that imputation quality decreases with increased African ancestry.

Lastly, SHAPEIT-IMPUTE2 with 1000G reference panel was successful in identifying G206A mutation carriers. We also noticed that imputation quality drastically improved when imputation was conducted using large (5MB) chunk size as compared to small (1MB) chunks. This seems to contradict previous observation: Zhang et al (Zhang et al., 2011) studied the effect of window size on imputation in an African-Americans. They concluded that window size of 1MB could be considered acceptable. Possible explanations for these different results might be the more complex admixture of CH compare to AA (three-way vs. two-way admixed) and a more complex LD pattern for the G206A region. Ultimately, we recommend to consider a wider window size to achieve high-quality imputation in specific variants that fail under default settings.

This work has limitations. First, we could carry out the comparison between the two reference panels restricting the analyses to overlapping variants only, limiting our observation to a subset of the variants included in the 1000G panel. This is a result of the HRC composition, which is composed by several studies and ended up including only a consensus number of variants. Second, we tested the agreement between imputed and sequenced variants in a smaller subset of individuals that had both GWAS and WES data available.

## Acknowledgments

We thank the EFIGA study participants and the EFIGA research and support staff for their contributions to this study. This study was supported by funding from the National Institute on Aging [R21AG054832 (GT); 5R37AG015473 and RF1AG015473 (RM); R56 AG051876 and R01 AG058918 (JL)] and the BrightFocus Foundation (A2015633S (JL)).

