## Supplementary material for "Rare variants imputation in admixed populations: Comparison across reference panels and bioinformatics tools": suppl_file

### 1) Commands

Analyses were performed on Linux Debian Stretch 9.4 computing cluster with 125G RAM.

#### 1.1 **Phasing**

#### A) ***Eagle2***

```
eagle --allowRefAltSwap --vcfTarget=input.vcf.gz --geneticMapFile=genetic_map.txt \
--vcfOutFormat=v --vcfRef=reference.bcf \
--outPrefix=phased_VCF --chrom=number --numThreads=16 2>&1 | tee phased_vcf_CHRnumber.log
```

VCF output were then converted into .hap format for IMPUTE2 imputation using script from http://mathgen.stats.ox.ac.uk/impute/impute_v2.html as

```
perl vcf2impute_legend_haps.pl -vcf input.vcf -chr 21 -leghap impute_haps
```

#### B) ***SHAPEIT***

#### Strand check

```
shapeit -check -B input_bed -M input_mapfile --input-ref ref_hap_file ref_legend_file ref_sample_file --output-log output_log
```

#### Phasing

```
shapeit -B input_bed_file -M input_map_file \
--input-ref ref_hap_file ref_legend_file ref_sample_file \
--duohmm --output-graph out_phased-duohmm.graph \
--output-max out_bed_file-duohmm \
--thread 16 --seed 123456789 -W 5
```

### 1.2 **Imputation**

#### A) ***IMPUTE2***

```
impute2 -use_prephased_g -int start_pos end_pos \
-o out_impute.impute2 \
-m genetic_file -known_haps_g duo_haps_file -h ref_hap_file \
-l ref_legend_file -Ne 20000
```

#### B) ***MaCH-Admix***

#### I) **Extract SNPs within range using Plink** (v1.9)

```
plink --bfile input --extract range file_with_range --make-bed --out SNPs_rangePlink
```

#### II) **Create merlin file format using King** (v1.9)

```
king -b input.bed --merlin --prefix input_admix
```

#### III) **MaCH-Admix** (v2.0.203)

```
mach-admix -d file.dat -p file.ped -h ref_vcf  --vcfRef --outputstart start_pos \
--outputend end_pos --autoflip --startposition flank_start_pos \
--endposition flank_end_pos --vcfDupMarkerAppendLineNo --geno --quality --dosage --uncompressed --prefix
output
```

#### 1.3 VCF compare

A) **Convert imputed genotype probabilities into plink format**

```
plink --gen post_prob.gen --hard-call-threshold 0.1 --sample input_sample.sample  --make-bed --out output_plink
```

B) **Extract SNPs**

```
vcftools --gzvcf input.vcf.gz --snps snp_list --recode --recode-INFO-all  --out output_vcf
bgzip -c output_vcf.recode.vcf > output_vcf.recode.vcf.gz
tabix -p vcf  output_vcf.recode.vcf.gz
```

C) **Compare**

```
vcf-compare -g sequenced.recode.vcf.gz reference.recode.vcf.gz  | grep ^GC  | cut -f 2-7,20-24 > vcf_compare
```

**Tools version**:

zlib: 1.2.8

bzip: 1.8-32-g9cd85ab

tabix: 1.8-32-g9cd85ab

1.4 **Wilcoxon Signed-Rank Test**

Wilcoxon Signed-Rank Test was performed in R as:

```
wilcox.test(value_a,value_b,paired=TRUE)
```

**1.5 Poisson regression**

```
library(sandwich)

summary(robust<-glm(SNP_Mismatches ~ Ancestry_Name, family="poisson", data=
data_frame))

cov.robust <- vcovHC(robust, type="HC0")

std.err <- sqrt(diag(cov. robust))

r.est <- cbind(Estimate= coef(robust), "Robust SE" = std.err,

 "Pr(>|z|)" = 2 * pnorm(abs(coef(robust)/std.err),

 lower.tail=FALSE), LL = coef(robust) - 1.96 * std.err,

 UL = coef(robust) + 1.96 * std.err)

print(r.est_yri)
```

### 2 Comparison of Phasing Tools: Eagle2 vs. SHAPEIT2

We compared counts between Eagle2-IMPUTE2 and SHAEPIT-IMPUTE2 for bi-allelic SNPs only. Initially, we assessed the percentage of “high-quality” variants regardless of their MAF. The percentage of “high-quality” SNPs was similar between phasing tools using the 1000G reference panel. However, when HRC panel was employed we observed a higher percentage of “high-quality” SNPs using SHAPEIT-IMPUTE2 as compared to Eagle2-IMPUTE2 (48.57% vs. 45.94%) (**Supplementary Table 1**). With 1000G reference panel, we observed comparable percentage of “high-quality” uncommon and ultra-rare SNPs for Eagle2 and SHAPEIT2 phasing tools except rare SNPs (**Supplementary Table 2**). Using HRC panel, we found that for “high-quality” SNPs SHAPEIT-IMPUTE2 retrieved 91.09% and 60.12% of uncommon and rare variants respectively as compared to 87.27% and 53.16% from Eagle2-IMPUTE2 (**Supplementary Table 2**). When we compared computational time between the phasing tools, we observed SHAPEIT2 was at least 5-times slower than Eagle2 (**Supplementary Table 3**).

#### Imputation with Eagle2 Phased data

For phasing with Eagle2 (v2.4), reference genome were downloaded from ftp://ftp.1000genomes.ebi.ac.uk/vol1/ftp/release/20130502/ Data were QC-ed and processed as per guidelines on https://data.broadinstitute.org/alkesgroup/Eagle/#x1-310005.3.1

### 3 Agreement

We used vcf-compare to compare genotypes as defined in workflow (**Supplementary Figure 1)**. In summary, overlapping SNPs based on chromosome, position, reference, alternate allele with HRC, 1000G imputed SNPs, and WES data were initially filtered. Next, SNPs with 0% missingness in WES data were obtained. For these SNPs for each reference panel, SNPs were filtered based on imputed info value, MAF from IMPUTE2 info files. Filtered SNPs were used to create VCF files for reference panel and WES data. Finally, vcf-compare utility was used to compare VCFs.

Mismatch percent was calculated as (Total count of Mismatch)/(Total count of Mismatch + Total count of match).

Total SNPs in VCF compare = (Number of individuals)*(SNPs). For example, for 262 individuals with 9,170 ultra-rare high-quality SNPs, total SNPs in VCF compare are 2,402,540.

Slope for individual ancestry was calculated in R (**Supplementary** section **1.5**)

### 4 **Cohen kappa statistic**

Cohen's kappa coefficient (*K*) was calculated as defined in workflow (**supplementary Figure 2)**. In brief, overlapping SNPs based on chromosome, position, reference and alternate allele with HRC, 1000G imputed SNPs and WES data were initially filtered. Next, SNPs with 0% missingness in WES data were obtained. For these SNPs for each reference panel, SNPs were filtered based on imputed info value, MAF from IMPUTE2 info files. Filtered SNPs were used to create .raw files using plink for reference panel and WES data. Finally, Cohen's Kappa (*K*) coefficient was calculated on filtered SNPs.

### 5 Ancestral effect on SNP mismatch

We used sandwich_2.5-0 R package to calculate SE error and P-values when Poisson regression was conducted. Slope for individual ancestry in different MAF bins was calculated in R (**Supplementary** section **1.5**)

**Supplementary Table 1** Comparison of Eagle2 and SHAPEIT2 phasing tools in tandem with IMPUTE2 for imputation on CHR21 using HRC and 1000G reference panels for input SNP counts and Bi-allelic SNPs based on different info threshold.

| **1000G** | | | | | |
|---|---|---|---|---|---|
| **Tool** | **Input Reference SNPs** | **Imputed SNPs (%)** | **Bi-allelic** | | |
| | | | **Total** | **Info ≥ 0.40 (%)** | **Info ≥ 0.80 (%)** |
| SHAPEIT - IMPUTE2 | 1,104,648 | 1,104,648 (100) | 1,055,422 | 446,591 (42.31) | 315,688 (29.91) |
| Eagle2 - IMPUTE2 | 1,105,538 | 1,091,773 (98.75) | 1,047,297 | 446,207 (42.60) | 308,869 (29.49) |
| **HRC** | | | | | |
| SHAPEIT - IMPUTE2 | 531,276 | 530,361 (100) | 530,361 | 327,483 (61.74) | 257,616 (48.57) |
| Eagle2 - IMPUTE2 | 531,276 | 530,361 (100) | 530,361 | 333,721 (62.92) | 243,693 (45.94) |

**Supplementary Table 2** Comparison of Eagle2 and SHAPEIT2 phasing tools in tandem with IMPUTE2 for imputation on CHR21 using HRC and 1000G reference panels in different Info threshold stratified by MAF bins.

| MAF | Eagle2-IMPUTE2 | | | SHAPEIT-IMPUTE2 | | |
|---|---|---|---|---|---|---|
| | Info ≥0 | Info ≥0.40 (%) | Info ≥0.80 (%) | Info ≥0 | Info ≥0.40 (%) | Info ≥0.80 (%) |
| **1000G** | | | | | | |
| [1 - 5%] | 79,557 | 78,880 (99.14) | 68,820 (86.50) | 80,960 | 80,332 (99.22) | 70,600 (87.20) |
| [0.1 - 1%) | 287,712 | 242,331(84.22) | 139,216 (48.38) | 280,289 | 240,890 (85.94) | 143,031 (51.02) |
| [0 – 0.1%) | 578,408 | 24,836 (4.29) | 3,790 (0.6552) | 591,417 | 24,058 (4.06) | 3,959 (0.6694) |
| **HRC** | | | | | | |
| [1 - 5%] | 69,101 | 68,884 (99.68) | 60,309 (87.27) | 73,691 | 73,446 (99.66) | 67,130 (91.09) |
| [0.1 - 1%) | 172,858 | 160,653 (92.93) | 91,892 (53.16) | 162,556 | 150,801 (92.76) | 97,733 (60.12) |
| [0 – 0.1%) | 197,395 | 13,346 (6.76) | 2,385 (1.20) | 202,527 | 11,818 (5.83) | 2,646 (1.30) |

**Supplementary Table 3** Comparison of phasing CPU times (seconds) in HRC and 1000G reference panel using SHAPEIT2 and Eagle2

| Reference Panel | Tool | |
|---|---|---|
| | **SHAPEIT2** | **Eagle2** |
| HRC | 27142.74 | 4042.704 |
| 1000G | 12660.104 | 2521.74 |

**Supplementary Table 4** Comparison of SHAPEIT-IMPUTE2 and MaCH-Admix imputation tools on CHR21 using 1000G for input SNP counts and Bi-allelic SNPs based on platform-specific quality index.

<table>
<tr><th colspan="5">SHAPEIT-IMPUTE2</th></tr>
<tr><th rowspan="2">Input Reference SNPs</th><th rowspan="2">Imputed SNPs (%)</th><th colspan="3">Bi-allelic</th></tr>
<tr><th>Total SNPs</th><th>Info ≥ 0.4 (%)</th><th>Info ≥ 0.80 (%)</th></tr>
<tr><td>1,104,648</td><td>1,104,648 (100)</td><td>1,055,422</td><td>446,591 (42.31)</td><td>315,688 (29.91)</td></tr>
<tr><th colspan="5">MaCH-Admix</th></tr>
<tr><td rowspan="2">653,791</td><td rowspan="2">646,594 (98.89)</td><th colspan="2">Total SNPs</th><th>Rsq ≥ 0.3 (%)</th></tr>
<tr><td colspan="2">598,943</td><td>598,943 (92.63)</td></tr>
</table>

**Supplementary Table 5** Comparison of SHAPEIT-IMPUTE2 and MaCH-Admix imputation tools on CHR21 using 1000G based on different quality threshold respective to tool and MAF bins.

| MAF | MaCH-Admix | | SHAPEIT-IMPUTE2 | | |
|---|---|---|---|---|---|
| | Rsq ≥ 0 | Rsq ≥ 0.30 (%) | Info ≥ 0 | Info ≥ 0.40 (%) | Info ≥ 0.80 (%) |
| [1% - 5%] | 74,139 | 74,139 (100) | 80,960 | 80,332 (99.22) | 70,600 (87.20) |
| [0.1% - 1%) | 171,909 | 171,909 (100) | 280,289 | 240,890 (85.94) | 143,031 (51.02) |
| [0 – 0.1%) | 252,476 | 12,802 (5.07) | 591,417 | 24,058 (4.06) | 3,959 (0.6694) |

**Supplementary Table 6** Comparison of CPU compute time (seconds) for MaCH-Admix and SHAPEIT-IMPUTE2 on region of 1MB on CHR21

| Reference Panel | Tool | Start position | End position | Time |
|---|---|---|---|---|
| 1000G | MaCH-Admix | 10,411,245 | 11,411,244 | 36492.78 |
| 1000G | SHAPEIT-IMPUTE2 | 10,000,000 | 10,999,999 | 333.744 |

**Supplementary Table 7** Comparison on effect of ancestral components for high-quality (IMPUTE2 info ≥0.80) SNPs for HRC and 1000G reference panels in different MAF bins.

| MAF | Ancestry | 1000G | | | | HRC | | | |
|---|---|---|---|---|---|---|---|---|---|
| | | **Estimate** | **P-value** | **Robust SE** | **Conf. Int.** | **Estimate** | **P-value** | **Robust SE** | **Conf. Int.** |
| [1% - 5%] | CEU | -0.16 | 2.17E-04 | 0.04 | [-0.24, -0.07] | -0.35 | 1.95E-08 | -0.35 | [-0.48, -0.23] |
| | NAT | -0.16 | 2.20E-05 | 0.03 | [-0.24, -0.09] | -0.20 | 0.037 | 0.09 | [-0.391, -0.01] |
| | YRI | 0.23 | 1.69E-09 | 0.03 | [0.15,0.30] | 0.40 | 5.66E-13 | 0.05 | [0.29, 0.52] |
| [0.1% - 1%) | CEU | -1.02 | 2.11E-05 | 0.24 | [-1.49, -0.55] | -1.14 | 3.12E-09 | 0.19 | [-1.52, -0.76] |
| | NAT | -1.63 | 2.99E-05 | 0.39 | [-2.40 -0.86] | -1.51 | 3.30E-06 | 0.32 | [-2.15, -0.87] |
| | YRI | 1.46 | 1E-15 | 0.18 | [1.10, 1.82] | 1.52 | 4.44E-24 | 0.15 | [1.23, 1.82] |
| [0 – 0.1%) | CEU | -0.50 | 0.72 | 1.40 | [-3.26, 2.25] | -0.64 | 0.42 | 0.81 | [-2.24, 0.94] |
| | NAT | -5.78 | 0.028 | 2.63 | [-10.94, -0.61] | 0.63 | 0.48 | 0.91 | [-1.16, 2.44] |
| | YRI | 1.58 | 0.16 | 1.13 | [-0.64, 3.81] | 0.11 | 0.88 | 0.79 | [-1.44, 1.66] |

**Supplementary Figure 1**

VCF compare workflow to obtain counts for mismatched and matched genotypes

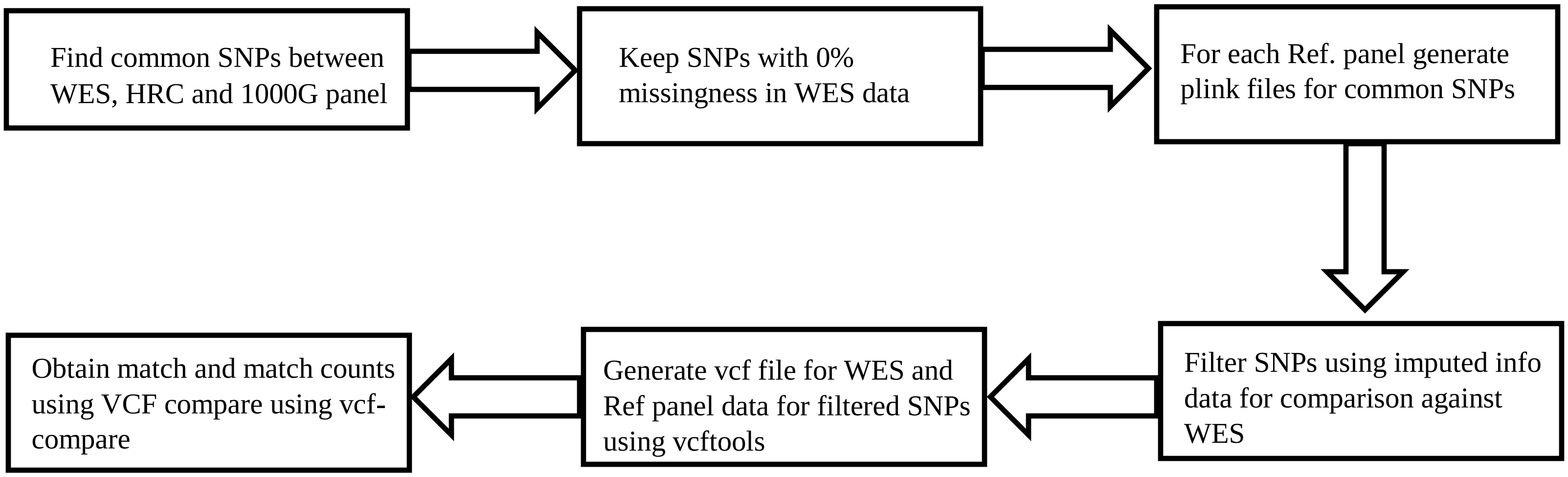

**Supplementary Figure 2**

Cohen's Kappa coefficient workflow

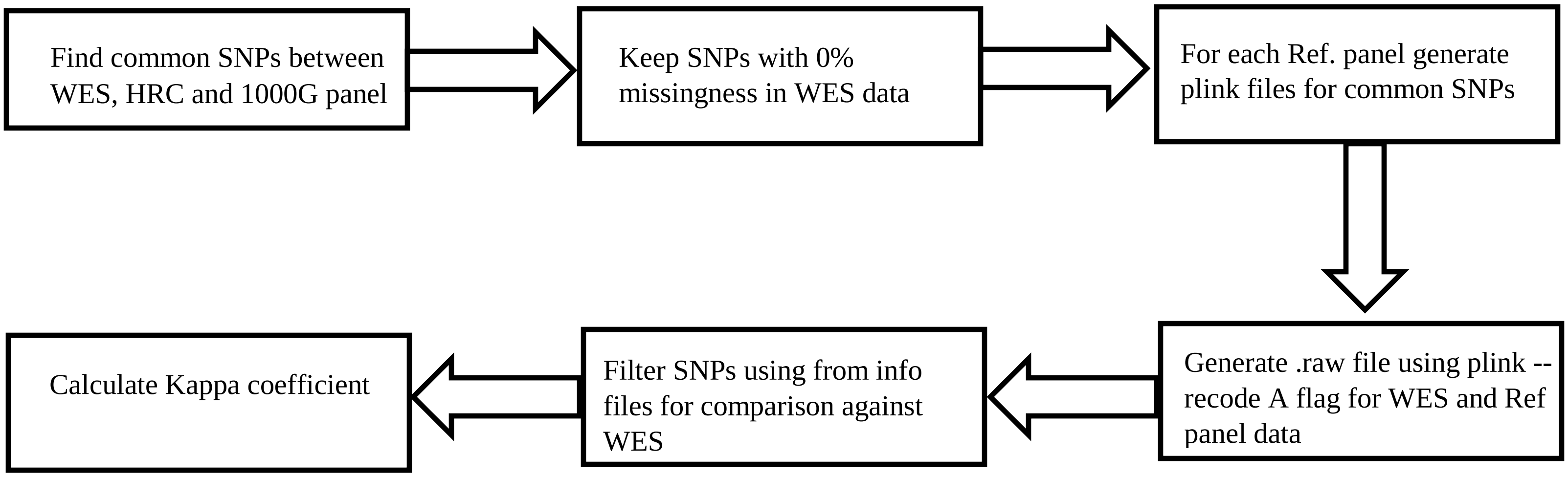
